# Depression in Parkinson’s disease is associated with dopamine unresponsive reduced reward sensitivity during effort-based decision making

**DOI:** 10.1101/2024.05.09.592897

**Authors:** Harry Costello, Yumeya Yamamori, Karel Kieslich, Mackenzie Murphy, Kamilla Bobyreva, Anette-Eleonore Schrag, Robert Howard, Jonathan P Roiser

## Abstract

Willingness to exert effort for a given goal is dependent on the magnitude of the potential rewards and effort costs of an action. Such effort-based decision making is an essential component of motivation, in which the dopaminergic system plays a key role. Depression in Parkinson’s disease (PD) is common, disabling and has poor outcomes. Motivational symptoms such as apathy and anhedonia, are prominent in PD depression and related to dopaminergic loss. We hypothesised that dopamine-dependent disruption in effort-based decision-making contributes to depression in PD.

In the present study, an effort-based decision-making task was administered to 62 patients with PD, with and without depression, ON and OFF their dopaminergic medication across two sessions, as well as to 34 patients with depression and 29 matched controls on a single occasion. During the task, on each trial, participants decided whether to accept or reject offers of different levels of monetary reward in return for exerting varying levels of physical effort via grip force, measured using individually calibrated dynamometers. The primary outcome variable was choice (accept/decline offer), analysed using both logistic mixed-effects modelling and a computational model which dissected the individual contributions of reward and effort on depression and dopamine state in PD.

We found PD depression was characterised by lower acceptance of offers, driven by markedly lower incentivisation by reward (reward sensitivity), compared to all other groups. Within-subjects analysis of the effect of dopamine medication revealed that, although dopamine treatment improves reward sensitivity in non-depressed PD patients, this therapeutic effect is not present in PD patients with depression.

These findings suggest that disrupted effort-based decision-making, unresponsive to dopamine, contributes to PD depression. This highlights reward sensitivity as a key mechanism and treatment target for PD depression that potentially requires non-dopaminergic therapies.

## Introduction

Depression in Parkinson’s disease (PD) is associated with greater disability^1^, increased mortality^2^and a greater negative impact on health related quality of life than motor symptoms^3^. Often occurring as an early symptom prodrome, at least one-third of people living with PD develop depression; consequently, effective treatment of depression in PD has the potential to achieve significant health and economic benefits.^4^ However, depression in PD goes undiagnosed in up to half of patients, and current treatments for depressed mood are poorly effective^5–8^. Despite significant progress in our understanding of the aetiology of motor symptoms in PD, we know very little about the underlying mechanisms of depression in PD, posing a major barrier to developing effective treatments.

Depression is a multifactorial, heterogeneous, and aetiologically complex syndrome.^9,10^ There are at least 256 possible unique symptom profiles that meet DSM-V criteria for a diagnosis of major depressive disorder. ^11^ Though the reliability of different depression subtypes has not been established in the general population,^12^ the aetiology of PD and the associated depressive symptom profile suggests that depression in PD may be driven by more common and homogenous underlying mechanisms.^13^

Motivational symptoms such as apathy and anhedonia are particularly prominent in PD, affecting 40% and 46% of patients, respectively^14^. Human and animal studies have shown that mesolimbic dopaminergic transmission is crucial for motivated behaviour and reward processing.^15^ Loss of pre-synaptic dopaminergic projections to the striatum and reduced functional connectivity in this region are associated with increased apathy and anhedonia in PD.^16,17^ This may explain why mood changes in PD are frequently associated with motor fluctuations, or “ON/OFF” dopamine states^18^, and suggests depression in PD could be related to dopaminergic deficits which mediate specific neurocognitive processes.

Biases in specific neurocognitive processes and their interaction with socioenvironmental factors over time are thought to drive the emergence of depressive phenotypes.^19^ Recently, computational methods have been used to experimentally dissect the cognitive components driving disorders of motivation.^15^ A core process of motivated goal directed behaviour explains how the potential benefit or reward for performing an action is evaluated with respect to the amount of effort required to attain it. Disruption to this process, termed effort-based decision making for reward, is believed to underlie motivational syndromes and potentially depression in Parkinson’s disease, in which motivational symptoms are prominent.^15^

Measures of effort-based decision making utilise tasks that elicit effects of decision-making on the relative potential costs and benefits of an action.^15^ For example, participants are asked to make decisions as to whether to exert varying levels of effort (e.g. via grip force) for different magnitudes of reward. Studies in PD and other neurological disorders have shown that changes in how reward is evaluated against effort costs contribute to apathy.^20–22^ For example, apathetic PD patients are less willing to exert effort for low rewards than more motivated patients.^20,23^

The mesolimbic dopamine system has been consistently implicated as a key neuromodulator of motivation and effort-based decision making.^24^ Depletion of ventral striatal dopamine in animal models shifts choice behaviour to selection of low-effort options.^25^ When OFF dopaminergic medication PD patients also exhibit greater impairments in reward processing.^26^ Specifically, dopaminergic medication has a general effect in motivating patient behaviour for high-effort, high-reward options.^20^ This may explain why treatment with dopamine agonists are associated with reduced motivational symptoms of depression over time in PD.^27^

Considering the effect of dopamine depletion on mood in PD, the high prevalence of motivational symptoms and existing evidence of disruption in effort-based decision making, we hypothesised that dopamine-dependent impairment in effort-based decision making is a key mechanism underlying PD depression. Specifically, we predicted that PD patients with depression would exhibit a combination of both lower reward sensitivity and greater effort sensitivity. We also predicted that PD patients would exhibit greater discounting of reward by effort (i.e. have greater effort sensitivity), when OFF dopaminergic medication, and that the effect of dopaminergic medication withdrawal would be greater in patients with depression and PD. All study predictions were preregistered after data collection had commenced but prior to any analysis (https://osf.io/b2umh). Please note that the preregistered sample size appears incorrect due to an initial duplication error during data entry; no participants were excluded.

To test this, we conducted the first study to investigate the effects of dopamine and depression on effort-based decision making in PD.

## Materials and methods

### Ethical approval

The study was approved by the local ethics committee (approved by HRA and Health and Care Research Wales, REC reference: 22/NS/0007, date of approval: 1^st^ February 2022), and written consent was obtained from all subjects in accordance with the Declaration of Helsinki.

### Participants

In this case-control study, four groups of age-(over 50 years old) and sex-matched participants were recruited (n=125): 29 healthy participants with no neurological or psychiatric conditions, 34 patients with major depressive disorder, 31 patients with Parkinson’s disease and 31 patients with Parkinson’s disease and major depressive disorder.

All PD patients had a clinical diagnosis of idiopathic PD confirmed by a neurologist and were recruited from movement disorders clinics in the Greater London area, UK. All participants with major depressive disorder were clinically assessed and diagnosed by a psychiatrist (H.C) according to DSM-V criteria. Exclusion criteria included evidence of dementia or other major neurological or psychiatric conditions (other than PD, depression or co-morbid anxiety), and use of dopamine modulating medications other than those indicated for PD (for example, antipsychotics or stimulants). To screen for other psychiatric conditions, all participants underwent the Mini-International Neuropsychiatric Interview (MINI), a validated short structured interview.^28^ As a baseline cognitive screen for dementia, all subjects were administered the Mini-Mental State Examination and excluded if they scored less than 26.^29^

Healthy control participants and depressed patients without PD underwent testing once. Both PD patient groups were tested twice in morning sessions, once ON, following their normal dopaminergic medications, and once having withheld their dopaminergic medication since the night before for at least ten hours prior to testing (OFF). These sessions were counterbalanced in order across both PD groups. A minimum of one week between testing sessions was ensured to minimise repetition effects.

### Clinical measures

Depression symptom severity was measured using the interviewer-rated 17-item Hamilton Depression Rating Scale (HAM-D)^30^ (score range: 0-50) and 21-item patient-rated Beck Depression Inventory (BDI)^31^ (score range: 0-63), both of which were selected as they are valid rating scales for depression in PD.^32^ Motivational symptoms were measured using the patient-rated Dimensional Anhedonia Rating Scale (DARS)^33^ and the 18-item clinician and patient-rated versions of the Apathy evaluation scale.^34^

Severity of PD was assessed using the Movement Disorder Society-Unified Parkinson’s Disease Rating Scale (MDS-UPDRS)^35^ and the Hoehn and Yahr stage.^36^

Participants also completed the Wechsler Test of Adult Reading (WTAR)^37^ as a proxy measure of intelligence quotient (IQ) and education.

All symptom severity measures were repeated in the ON and OFF states for PD participants.

Demographics and baseline measures for all groups are reported in Table 1.

### Effort-based decision-making task

Participants completed an effort-based decision-making task, the ‘apple-gathering task’, which has previously been used in patients with PD^20,38^. This paradigm was designed in Psychtoolbox (psychtoolbox.org) within MATLAB and administered on a laptop computer.

On each trial, participants were asked to either accept or reject offers of different levels of reward in return for exerting different levels of physical effort (grip force) using their non-dominant hand (see Figure 1). The key outcome measure was willingness to accept or reject a challenge (the decision phase). Participants registered their responses using arrows on the keyboard (left for ‘yes’, right for ‘no’) and a handheld dynamometer (SS25LA, BIOPAC Systems), to capture effort exertion via grip force. Each participant’s maximum voluntary contraction (MVC) was calculated as the greatest force exerted over three maximal contractions and was established for each testing session so that force levels were calibrated and thus normalised on an individual participant basis.

**Figure 1.**
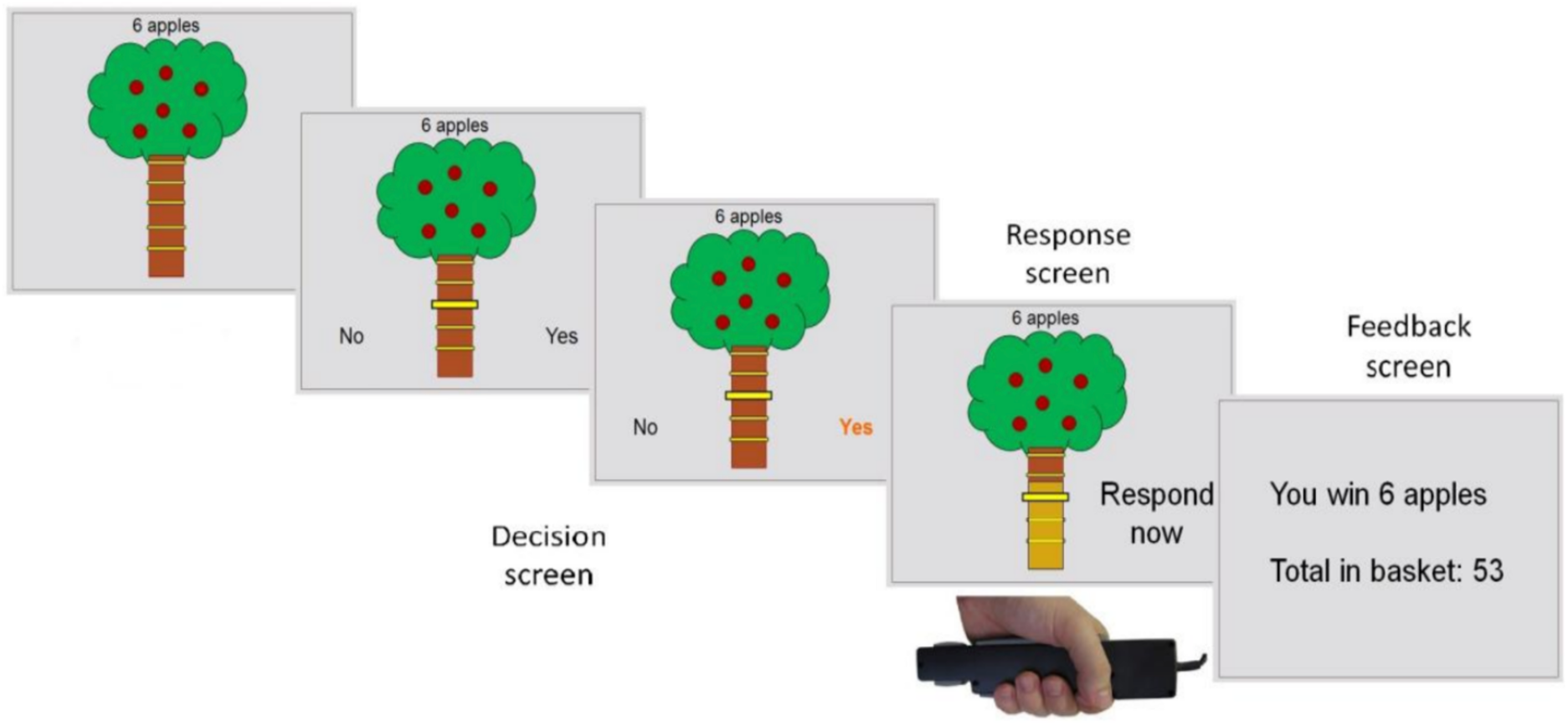
The Apple Gathering Task (AGT) In this task participants decide to accept or reject offers within which different levels of reward are available for different levels of physical effort (grip force) (A). The key outcome measures are the willingness to accept a challenge (the decision phase). Each participant’s maximum voluntary contraction (MVC) is established so that force levels are calibrated on an individual basis. Participants are asked to squeeze the handles of the dynamometer as hard as they can with their right and left hands. Participants are instructed to gather as many apples as they choose to over the course of the experiment, knowing that the money they receive at the end depends on the total number of apples gathered. The image on screen displays the reward on offer (number of apples on a tree) and the force required to obtain that level of reward (height of the yellow line on the tree trunk). There are four different reward levels (3, 6, 9, 12) and four effort levels 20/40/60/80% MVC.

Each offer was presented on the screen as an apple tree (see Figure 1). The reward on offer for each trial was indicated by the number of apples on the tree (3, 6, 9 and 12 apples), while effort required was indicated by the height of a yellow line on the tree trunk. Effort levels corresponded to 20, 40, 60, and 80% of the participant’s MVC. Participants were offered monetary rewards proportional to the total number of apples they collected. The four reward and four effort levels were orthogonally combined (to give 16 conditions) and presented in a pseudo-randomised order across five blocks, totalling 80 trials. All participants received the same offers presented in the same order.

Prior to starting the experiment, participants practised each effort level to familiarise themselves with the force required and completed a practice block consisting of four combinations of effort and reward.

Participants were instructed to weigh up the effort costs against the reward offered for each trial, and to decide ‘whether it is worth it’. If they accepted an offer (by exerting a small squeeze on the dynanometer) they then had to squeeze to the required force and hold this for >1s before being rewarded with the apples. Prior to starting the experiment, participants practised each effort level to familiarise themselves with the force required and completed a practice block consisting of four combinations of effort and reward.

### Model-agnostic analyses

Model-agnostic analyses were conducted in R. To account for the possibility that probabilistic discounting drives decision making in the task (i.e. participants not accepting higher effort offers due to physically being unable to exert the effort required, despite having passed the calibration phase of the task), we initially analysed the effect of success rate on offer acceptance. As success rates were positively associated with acceptance rates, we covaried for success rate in all mixed models described below in sensitivity analyses.

#### Mixed-effects regressions of trial-by-trial choices

##### Case-control analysis

Given the hierarchical nature of our study, generalised logistic mixed effects models were used to model accept/reject responses. We included a random effect of subject, reward and effort (mean corrected), and fixed effects of reward, effort and group. Mixed effects modelling comparing group effects across all four groups included responses from PD participants when tested in the ON-dopamine state.

A three-way interaction term between all fixed effects was modelled, as well as all two-way interactions and main effects.

##### Dopamine state analysis

Within-subjects modelling of the effect of dopamine state on task, which included only groups with Parkinson’s disease, also utilised a generalized linear mixed effects model with a logistic link function to model accept/reject responses. Mixed effects modelling included a random effect of effect of subject, reward and effort (mean corrected), in addition to fixed effects of reward, effort, dopamine medication state and depression group. A four-way interaction term between all fixed effects was modelled, as well as all three-way and two-way interactions and main effects.

Exploratory analysis using linear mixed effects modelling was also used to explore associations between depressive symptom severity, anhedonia and apathy and overall willingness to accept offers, and with effort and reward sensitivity, and how these symptoms interacted with dopamine medication state in PD patients.

Mixed model fit was tested using the Akaike information criterion (AIC) which penalises models for complexity, to counter over-fitting.

### Hierarchical Bayesian computational modelling

To parse the relative contributions of different cognitive processes that influence decision making during the task, a hierarchical Bayesian computational model incorporating a logistic link function was constructed such that we could model decisions based on the effort and reward level of a given trial. This allowed comparison of competing hypotheses of reward and effort contributions to decision making, in addition to estimating parameters corresponding to reward and effort processing for each individual, enabling group comparison and to examine associations with symptom scores.

To reduce the risk of type-I error, all participants were fitted under the same group-level priors. All models were implemented using the probabilistic modelling language Stan and parameters were estimated using Hamiltonian Markov Chain Monte Carlo (MCMC) sampling using soft constraints on likely parameter prior distributions, assuming participants come from a single group-level distribution. Twenty MCMC chains were run, each having 5000 samples.

A variety of model iterations with varying complexity were built to capture the contribution of various cognitive processes (supplemental Figures S4-6). For example, model iteration included the addition of quadratic terms for effort and reward parameters, fixed effort and reward parameters, and a stochasticity or ‘guess’ parameter. Once all models were fitted to the data, model comparison was performed to select the winning model using difference in ELPD (Expected Log Pointwise Predictive Density); a measure of the difference in expected log likelihood of new data points under the different models. This quantifies the difference in how accurately models predict new data with the lowest score identifying the most parsimonious model.

For brevity, we only report the winning model in the main text (see supplement for other models), which was the simplest model comprising an effort sensitivity term (linear), reward sensitivity term (linear) and an accept bias term. The model works as follows on a given trial:

i. Effort sensitivity – effort level is transformed through a linear effort sensitivity parameter to yield a subjective value of effort (the more negative, the more effort sensitive):

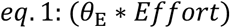

ii. Reward sensitivity – reward level is transformed through a linear reward sensitivity to yield a subjective value of the magnitude of reward such that a reward sensitivity <1 results in rewards being perceived as less rewarding than they truly are:

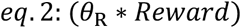

iii. Subjective value: Reward and effort parameters are then combined to form the subjective value of the offer:

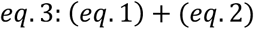

iv. Accept bias: the subjective value of the offer is passed through an invert logit link function with a bias parameter for each participant which maps the subjective value of the offer to a probability of accepting the offer which shifts the curve by a constant. This term therefore represents the tendency of accepting an offer independent of reward or effort level (the higher the bias term, the more likely a participant to accept offers):

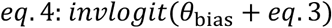

The same winning model was used for modelling of dopamine state effects on decision making. To capture the within-subjects design, we modelled the natural pairings of each parameter across conditions (i.e. the reward sensitivity ON and OFF dopaminergic medication) by drawing them from multivariate, correlated distributions. This approach has recently been shown to improve parameter estimation by allowing for the pooling of data across conditions, leading to within-parameter shrinkage.^39^

Convergence checks were conducted by visualizing trace plots and computing R-hat statistics across MCMC chains. Posterior predictive checks were conducted to ensure model predictions could accurately retrieve behavioural patterns in the original dataset.

## Results

Participant characteristics are presented in Table 1. The groups differed significantly on standardised WTAR (higher in healthy controls relative to all other groups) and age (lower in depressed relative to all other groups), and therefore these were included as covariates in all case-control models. As expected, duration since first onset of depressive symptoms was longer in the depressed (mean=27.2 years, sd=11.3) than the PD depressed (mean=12.9 years, sd=17.5) group. Other than this, the groups did not differ significantly on any other variable including PD patient motor symptom severity, total daily levodopa equivalent dose (LED), minutes since last dose prior to tests or change in motor symptom score ON and OFF dopamine. A total of five patients (PD group n=3, PD depressed group n=2) were unable to tolerate testing OFF dopaminergic medication and as a result, were only tested in the ON state.

We found that the addition of random effects of reward and effort, relative to random effects of subject alone, improved model evidence (AIC 5346 vs 4539), and we therefore subsequently used this as the model for all primary analyses. However, the addition of random effects that are also incorporated as fixed effects significantly reduces statistical power and increases variance, and therefore for completeness we additionally report the model with fixed effects of reward and effort (retaining random effects of subject: see supplement). As expected, in the model treating reward and effort as random effects, across all groups acceptance rates on the AGT increased significantly as reward levels increased (odds ratio (OR)= 1.76, 95%CI(1.43 to 2.15), p<0.001) (see Figure 2A and supplemental Figure S1) and decreased significantly as effort levels increased (OR=0.00, 95%CI(0.00 to 0.00), p<0.001) (see Figure 2B and supplemental Figure S1). No significant reward-by-effort interaction was found in the model treating reward and effort as random effects (OR=1.12, 95%CI(0.84 to 1.48), p=0.4), but this interaction was significant in the model in which they were incorporated as fixed effects (OR=1.48, 95%CI(1.21 to 1.81), p<0.001) (see Figure 2 and supplemental Figure S1). Post-hoc analysis showed that while higher effort offers resulted in significantly lower offer acceptance at all reward levels, this pattern was more pronounced at lower reward levels (contrast between lowest and highest effort levels at the lowest reward level: logOR=5.03, p<0.001) compared to higher reward levels (contrast between lowest and highest effort levels at the highest reward level: logOR=2.78, p< 0.001). In other words, effort has a progressively smaller effect on acceptance as reward levels increase.

**Figure 2.**
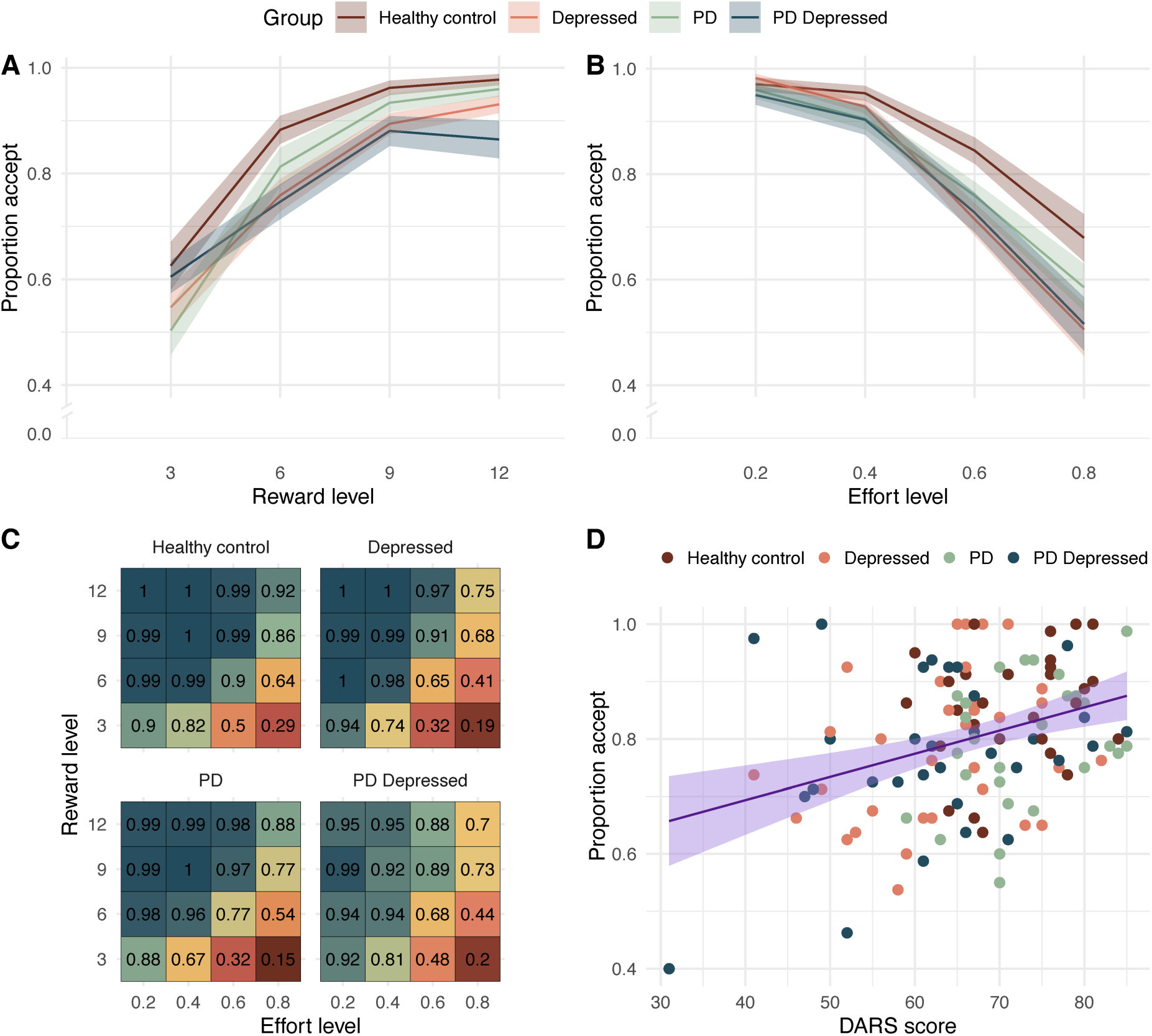
Empirical data showing a dissociation in acceptance between groups as effort, reward and symptom severity increased. **A.** Empirical data showing increase in acceptance as offer reward level increases and divergence in acceptance between groups (mean ± standard error). **B.** Empirical data showing reduction in acceptance as offer effort level increases and divergence in acceptance between groups (mean ± standard error). **C.** Change in each group acceptance as both reward and effort increases (mean change). **D.** Decrease in offer acceptance, irrespective of effort or reward, as anhedonia increases (lower DARS score)

To examine whether effort-based decision making was influenced by success rate, especially on high-effort trials (which could potentially induce probabilistic discounting), we also analysed success rates during effort exertion (only on accepted trials) (see supplemental Figure S2). Success rates decreased significantly as effort level increased (odds ratio (OR)=0.00, 95%CI(0.00-0.01), p<0.001). Importantly, however, there was no significant effect of group on success rate, or interactions between group and effort level (effort-by-group interaction, healthy volunteers relative to: depressed, OR=0.2, 95%CI(0.00 to 33.9), p=0.5; PD, OR=0.03 95%CI(0.00 to 12.3), p=0.3; PD depressed, OR=1.85, 95%CI(0.02 to 158), p=0.8). Nonetheless, we included individual mean success rate at each effort level in all subsequent models as this slightly improved model fit (AIC with covarying for success rate vs without: 5346 vs 5354).

### Depression in Parkinson’s disease is associated with weakened influence of reward on decisions

#### Mixed effects modelling of trial-level choices

We first assessed the effects of depression and PD on choice by comparing group performance in the ON dopamine state. There was a significant main effect of group on proportion of offers accepted. Depressed PD patients accepted significantly fewer offers than the PD and healthy control groups (main effect of group, compared to depressed PD ON: healthy controls, OR=17.8, 95%CI(3.83 to 83.0), p<0.001; PD ON, OR=5.13, 95%CI(1.34 to 19.7), p=0.017). Depressed patients without PD also accepted significantly fewer offers relative to healthy controls (main effect of group, compared to depressed: healthy controls, OR=8.25, 95%CI(1.71 to 39.7), p=0.008). Acceptance of offers was also lower in the PD depressed compared to the depressed group in models treating reward and effort as fixed effects (main effect of group, compared to depressed PD ON: depressed OR=4.23, 95%CI(1.66 to 10.8), p=0.003). However, this finding did not remain significant when treating reward and effort as random effects (main effect of group, compared to PD ON: depressed, OR=2.01, 95%CI(0.55 to 7.41), p=0.3).

Logistic mixed-effects modelling examining how reward and effort levels altered decision making revealed markedly lower incentivisation by reward in depressed PD patients. Depressed PD patients were less likely to accept offers as reward increased compared to all other groups (reward-by-group interaction, depressed PD ON relative to: healthy controls, OR=1.84, 95%CI(1.30 to 2.60), p<0.001; depressed, OR=1.49, 95%CI(1.11 to 2.00), p=0.007; PD ON, OR=1.73, 95%CI(1.27 to 2.35), p<0.001; Figure 2A). A comparison of all other groups revealed that this effect was specific to PD-depressed patients, as there was no significant impact of reward on offer acceptance between the other groups (reward-by-group interaction, group relative to healthy controls: depressed, OR=0.83, 95%CI[0.59 to 1.18], p=0.3; PD ON, OR = 0.97, 95% CI [0.67 to 1.39], p=0.9).

This pattern of results was similar when repeating the analysis with PD groups in the OFF dopamine state (reward-by-group interaction, group relative to PD depressed OFF: healthy controls, OR=1.56, 95%CI(1.18 to 2.07), p=0.002; PD OFF, OR=1.49, 95%CI(1.16 to 1.90), p=0.002). However, the reward-by-group interaction effect comparing the PD depressed OFF with the depressed group was no longer significant in the primary model incorporating random effects of reward and effort (OR=1.22, 95%CI(0.97 to 1.54), p=0.09).

We found no significant difference between groups in the effect of effort level on proportion of offers accepted when PD groups were in either ON (effort-by-group interaction, group compared to healthy controls: depressed, OR=0.12, 95%CI(0.00 to 12.7), p=0.4; PD ON, OR=1.33 95%CI(0.01 to 152), p>0.9; PD depressed ON, OR=4.01, 95%CI(0.04 to 393), p=0.6) or OFF dopamine state (effort-by-group interaction, group compared to healthy controls: depressed, OR=0.1, 95%CI(0.00 to 7.29), p=0.3; PD OFF, OR=0.12 95%CI(0.00 to 9.87), p=0.3; PD depressed OFF, OR=0.16, 95%CI(0.00 to 11.9), p=0.4).

Additionally, there was no significant three-way interaction between group, effort and reward in the primary model treating reward and effort as random effects when PD groups were in either the ON (group-by-effort-by-reward interaction, PD depressed ON compared to: healthy controls, OR=0.59, 95%CI(0.29 to 1.22), p=0.2; depressed, OR=0.64, 95%CI(0.40 to 1.04), p=0.07; PD ON, OR = 0.93, 95%CI(0.56 to 1.54), p=0.8), or OFF state (group-by-effort-by-reward interaction, PD depressed OFF compared to: healthy controls, OR=0.98, 95%CI(0.46 to 2.07), p>0.9; depressed, OR=1.11, 95%CI(0.51 to 2.42), p=0.8; PD OFF, OR=1.52, 95%CI(0.72 to 3.21), p=0.3).

To assess the influence of symptom severity beyond the group differences mentioned above, we modeled trial-by-trial offer acceptance, incorporating symptom scores (mean-corrected) as fixed effects in separate analyses, while controlling for group. This cross-group analysis included all healthy control, depressed, PD ON and PD depressed ON data. This analysis showed that participants with higher levels of anhedonia, depression, or apathy accepted fewer offers (DARS: OR = 1.07, 95%CI(1.02 to 1.12), p=0.004, BDI: OR = 0.91, 95%CI(0.86 to 0.97), p=0.004, AES-self: OR = 1.1, 95%CI(1.03 to 1.17, p=0.003) and were less incentivised by higher rewards levels (reward-by-symptom measure interaction, adjusting for group: DARS, OR=1.01, 95%CI(1.00 to 1.02), p=0.009; BDI, OR=0.98, 95%CI(0.97 to 1.00), p=0.012; AES-self, OR=1.02, 95%CI(1.01 to 1.03), p=0.006) (see supplemental Figure S3). To explore whether there was any pattern of trial-by-trial decision making which dissociated anhedonia from mood symptoms, mixed effects modelling was repeated for DARS score while controlling for BDI score and group. This analysis revealed that more severe anhedonia remained a significant predictor of lower offer acceptance (DARS: OR = 1.06, 95%CI(1.01 to 1.12), p=0.013). Furthermore, acceptance of offers increased less with increasing reward in individuals with more severe anhedonia symptoms (reward-by-DARS interaction: OR=1.01, 95%CI(1.00 to 1.02), p=0.009), over and above other mood symptoms or diagnostic group.

### Lack of willingness to exert effort in PD depression is driven by lower reward sensitivity

#### Computational modelling

As described in the methods, the best performing model was the simplest, incorporating: a linear reward parameter (reward sensitivity), a linear effort parameter (effort sensitivity) and an intercept parameter (accept bias), which accounts for the overall tendency to accept offers. Posterior predictive checks demonstrated that the winning model predictions were qualitatively similar to the raw data, recovered the pattern of behaviour observed in the model-agnostic analysis and almost perfectly recovered individual differences in acceptance rates (see supplemental Figures S4-6 for model comparison, and parameter recovery).

Modelling revealed a striking and specific difference between depressed PD patients relative to the other three groups in reward sensitivity (Figure 3). Depressed PD patients exhibited markedly lower reward sensitivity compared to all other groups (PD depressed group relative to: healthy controls, β=0.4, 95%CI(0.17 to 0.63), p<0.001; depressed, β=0.24, 95%CI(0.02 to 0.46), p=0.035; PD, β=0.45, 95%CI(0.23 to 0.67), p<0.001) (Figure 3). This suggests that during decision making depressed PD patients perceived potential rewards as less valuable. There were no significant differences between groups on accept bias and effort sensitivity.

**Figure 3.**
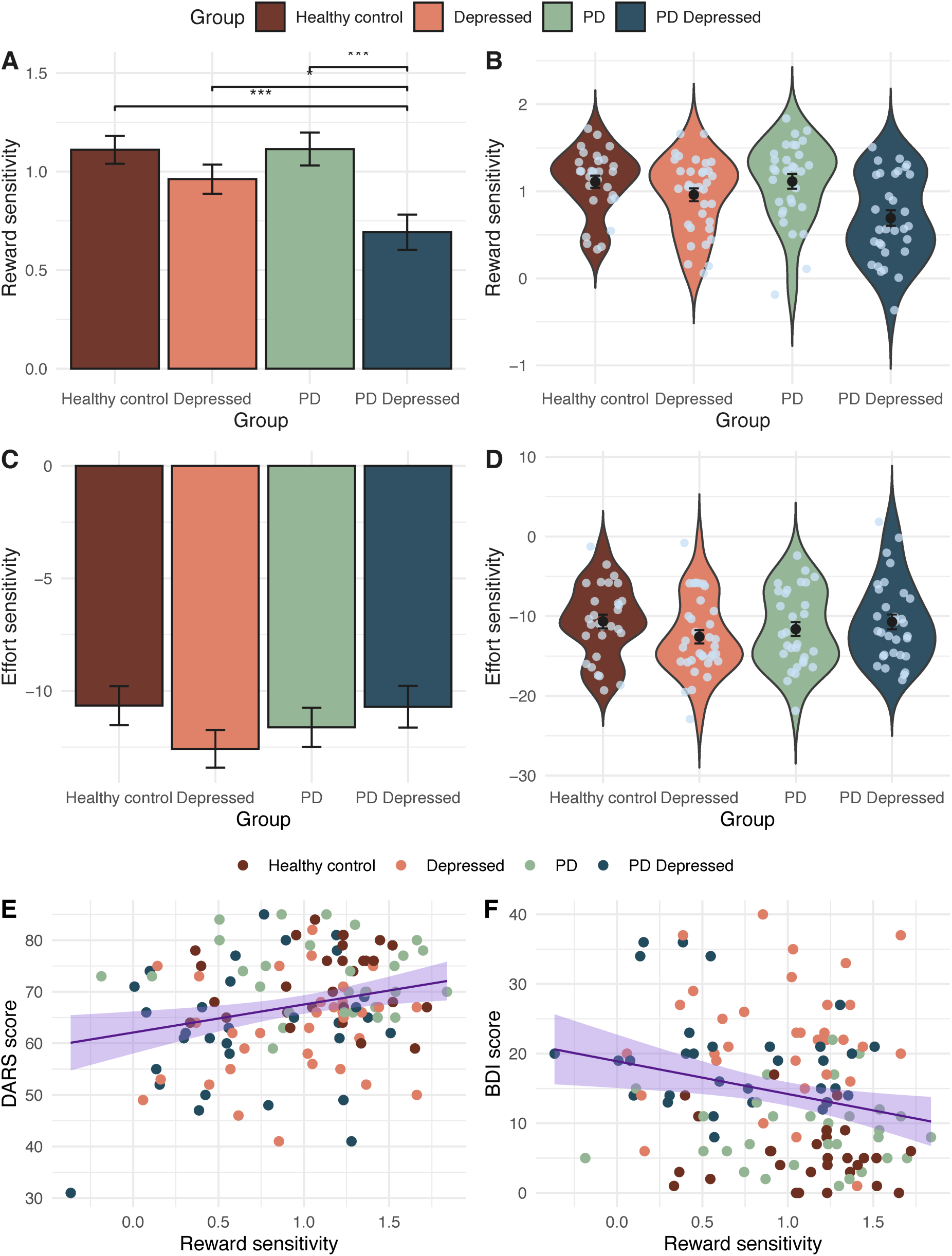
Depression in Parkinson’s disease is driven by reduced reward sensitivity. **A.** Depressed Parkinson’s patients exhibit significantly lower reward sensitivity than all other groups. **B.** Distribution of this difference in reward sensitivity and individual variation is shown in this group violin plot. In contrast there was no significant difference between groups in effort sensitivity (**C.** & **D.**). **E.** Across groups the more the anhedonic (lower DARS score) or (**F.**) depressed (higher BDI score) participants were the lower their sensitivity to reward. (reward sensitivity units: change in proportion accept with increase in stake available, effort sensitivity units: change in proportion accept with increase in effort required, ***p<0.001, **p<0.01, *p<0.05, bars and points in figures A-D represent parameter means, error bars = standard error)

In addition to the above group differences, more severe anhedonia (DARS: β=5.5, 95%CI(1.7 to 9.2), p=0.005) (Figure 4c), depression (BDI: β=-4.8, 95%CI(−8.2 to −1.3), p=0.007) (Figure 4d) and apathy (AES-self: β=5.1, 95%CI(2.1 to 8.2), p=0.001) symptoms were associated with lower reward sensitivity. However, when co-varying for group symptom severity, analyses were no longer significant, suggesting that this result is recapitulating group differences. Interestingly, however, when restricting this analysis to depressed participants (with and without PD) and covarying for group, anhedonia symptom severity narrowly missed significance (DARS: β=5.4, 95%CI(−0.41 to 11), p=0.068), suggesting specifically depressed patients with more severe anhedonia may subjectively perceive rewards as less valuable. To further investigate whether anhedonia was specifically associated with lower reward sensitivity, over and above other depressive symptoms, we repeated DARS analysis, across all groups, adjusting for BDI total score after removing BDI anhedonia items.^40^ The BDI anhedonia subscore has previously been validated and comprises the following items: loss of pleasure (item #4), loss of interest (item #12), loss of energy (item #15), and loss of interest in sex (item #21).^40^ This exploratory analysis revealed that lower reward sensitivity was significantly associated with greater anhedonia after adjusting for non-anhedonia depressive symptoms (DARS: β=3.6, 95%CI(0.04 to 7.2), p=0.047).

**Figure 4.**
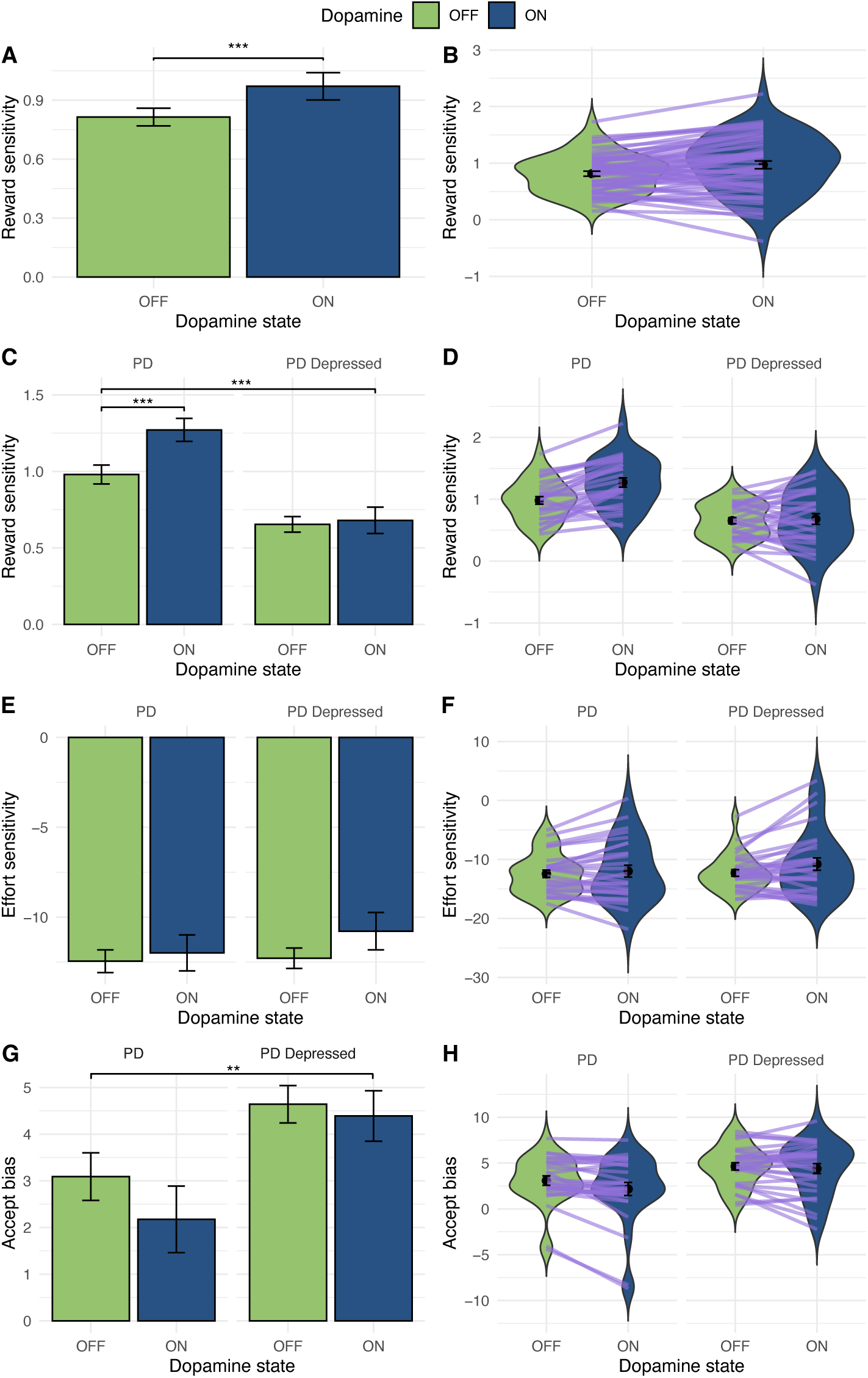
Dopamine treatment increases reward sensitivity in Parkinson’s disease but not Parkinson’s depression. Across all PD patients being ON dopamine increased reward sensitivity (**A** & **B**), purple lines represent individuals reward sensitivity change when ON and OFF dopamine). However, there was a dissociation between groups in dopamine-dependent incentivisation by rewards (**C** & **D**) where depressed PD patient reward sensitivity did not increase when ON dopamine compared to OFF. This dopamine-dependent dissociation between groups was not seen with effort sensitivity (**E** & **F**). Across both PD groups a significant reduction in accept bias when ON compared to OFF (**G** & **H)** dopamine was found but this did not remain significant after adjusting for reward sensitivity. (reward sensitivity units: change in proportion accept with increase in stake available, effort sensitivity units: change in proportion accept with increase in effort required, ***p<0.001, **p<0.01, *p<0.05, bars and points in all figures represent parameter means, error bars = standard error)

### Dopamine treatment increases reward sensitivity in Parkinson’s disease but not Parkinson’s depression

To determine the effect of dopamine mediation (ON vs OFF) on willingness to exert effort in PD, we used a within-subjects logistic mixed-effects model incorporating interaction terms, examining the effects of reward, effort, dopamine state and PD group. There was a significant two-way dopamine-by-PD group interaction (OR= 0.40, 95%CI(0.22 to 0.71), p=0.002) and three-way interaction between reward, dopamine and PD group (OR= 0.75, 95%CI(0.64 to 0.87), p<0.001); this indicates a dissociation in dopamine-dependent incentivisation by rewards between PD groups with and without depression (supplemental Figure S7). To understand this interaction, the effect of dopamine and the dopamine-by-reward reward interaction for each PD group were modelled separately. While PD patients were significantly more likely to accept offers overall when ON dopamine (relative to OFF dopamine: OR= 1.66, 95%CI(1.01 to 2.75), p=0.048), the opposite was the case for the PD depressed group (PD depressed ON relative to OFF dopamine: OR= 0.65, 95%CI(0.49 to 0.88), p=0.004). Furthermore, acceptance of offers increased less in the PD depressed group with increasing reward when ON relative to OFF dopamine medication (PD depressed reward-by-dopamine interaction: OR= 0.81, 95%CI(0.75 to 0.88), p=0<0.001). In contrast to reward, the equivalent three-way interaction between effort, dopamine and PD group was non-significant (OR= 9.23, 95%CI(0.86 to 99.3), p=0.067).

To help parse the factors driving this interaction, we performed within-subjects hierarchical Bayesian computational modelling, using the same winning model. This analysis revealed a robust effect of dopamine on reward sensitivity in PD patients without depression, but this effect was not present in the depressed PD group (PD group-by-dopamine state interaction: β=-0.27, 95%CI(−0.42 to −0.11), p<0.001) (see Figure 4). In other words, dopamine treatment increases reward sensitivity in PD patients (β=0.29, 95%CI(0.19 to 0.39), p<0.001), but this effect of treatment is not present in PD patients with depression (β=0.03, 95%CI(−0.09 to 0.15), p=0.7).

Supporting this finding, analysis of the interaction between dopamine state and symptom severity on reward sensitivity revealed that patients with more severe anhedonia (DARS: β=0.01, 95%CI(0.00 to 0.02), p=0.031) and depressive symptoms (BDI: β=-0.01, 95%CI(−0.02 to 0.00), p=0.029) had a lower response to dopaminergic medication, even after co-varying for group.

Similar to the above model-agnostic results, there was no main effect of dopaminergic medication on effort sensitivity (β=0.46, 95%CI(−0.63 to 1.6), p=0.4) or interaction with group (β=1, 95%CI(−0.50 to 2.6), p=0.2). Across both PD groups a significant reduction in accept bias when ON dopamine was found (main effect of dopamine medication: β=-0.92, 95%CI(−1.5 to −0.31), p=0.004), suggesting that dopaminergic treatment also reduces the likelihood of accepting offers in PD irrespective of potential rewards or effort costs. However, further analysis revealed that accept bias was significantly correlated with reward sensitivity (β=-0.06, 95%CI(−0.08 to −0.03), p<0.001). When analysis was repeated adjusting for reward sensitivity, there was no significant main effect of dopamine state on accept bias (main effect of dopamine medication on accept bias after adjusting for reward sensitivity: β=-0.45, 95%CI(−1.1 to −0.24), p=0.2). By contrast, the main effect of dopamine medication on reward sensitivity remained significant after adjusting for accept bias (main effect of dopamine medication on reward sensitivity after adjusting for accept bias: β=0.25, 95%CI(0.14 to 0.37), p<0.001), as did the PD group-by-dopamine state interaction (β=-0.24, 95%CI(−0.40 to −0.08), p=0.004).

To investigate whether the PD group-by-dopamine state interaction effect may be a consequence of the difference in antidepressant medication use between groups we repeated the analysis adjusting for antidepressant status. The main finding that reward sensitivity is dopamine-unresponsive in PD depression was unchanged (PD group-by-dopamine state interaction: β=-0.26, 95%CI(−0.42 to −0.10), p=0.002). Additionally, exploratory analysis exclusively within the PD depressed group but splitting participants by antidepressant status, found no association between use of antidepressant and reward sensitivity (β=0.04, 95%CI(−0.25 to 0.32), p=0.8) or any antidepressant-by-dopamine state interaction (β=0.00, 95%CI(−0.27 to 0.26), p>0.9).

## Discussion

This study demonstrates that depression in PD is associated with disrupted effort-based decision-making. Specifically, this effect is driven by reduced incentivisation by reward, rather than increased sensitivity to effort costs. Although dopamine treatment also impacts effort-based decision-making and increases reward sensitivity in PD, reward sensitivity in depressed PD patients is unresponsive to dopaminergic therapies. Our findings indicate that the disruption in reward sensitivity observed in depressed PD patients is specifically linked to motivational symptoms, particularly anhedonia, above and beyond other depressive symptoms such as dysphoric mood. This provides a clear plausible mechanism for the prominent motivational deficits and lack of interest in pleasurable activities that characterises depression in PD. The lack of response to dopaminergic medication may also explain why patients with depression exhibit a more persistent state of amotivation and depressed mood than the fluctuations in these symptoms described by PD patients without clinically significant depression who may have treatment responsive reward sensitivity deficits.^41^

Reward related signalling is crucial for goal-directed behaviour and disruption to this process has been associated with ‘decisional anhedonia’, where potential actions appear less rewarding leading to a cycle of amotivation, and loss of expected pleasure or reward from future actions.^42^ Impairment in reward valuation has been reported in depression without PD, although results have been inconsistent and a previous study using the same task did not find significant changes in reward sensitivity compared to healthy controls.^38,43^ In contrast, apathy in PD has been consistently associated with disrupted valuation of rewards, specifically low reward options.^20,23,44^ However, dopamine’s effect on effort-based decision making was the same in both apathetic and non-apathetic PD patients, increasing sensitivity to reward particularly for high effort options.^20^ In contrast our finding that depressed PD patients demonstrate a dopamine non-responsive reward sensitivity deficit, indicates a distinct mechanism underlying the effect of dopamine on depression and apathy in PD. Furthermore, our results underscore the value of using computational modelling to dissect latent drivers of behaviour that are not directly accessible through descriptive analyses alone. For instance, while our initial mixed-effects model suggested that PD depressed participants were more incentivised by reward when OFF compared to ON dopamine, this counterintuitive pattern was not evident when accounting for other latent cognitive processes through computational modelling. Instead, computational analysis revealed that reward sensitivity in PD depressed patients is unaffected by dopamine treatment. Instead, dopamine medication appears to change acceptance bias and effort sensitivity in PD depression, albeit the changes in the latter did not achieve statistical significance.

Optogenetic animal studies and human fMRI studies have identified a convergent network of brain regions involved in signalling reward valuation and effort costs, including the ventral striatum, anterior cingulate cortex (ACC) and orbitofrontal cortext (OFC).^15^ For example, optogenetic stimulation of dopaminergic projections from the ventral tegmental area increases reward seeking and striatal activity, whereas stimulation of the OFC reduces striatal response and reward seeking.^45^ In PD patients, reduced functional connectivity and striatal neurodegeneration precede the emergence of motivational symptoms.^16,17^ This suggests that disruption to frontostriatal circuit synchrony may impair reward valuation and lead to the emergence of motivational and mood symptoms. Different subregions of the ventral striatum have been shown to have dissociable contributions to the motivational versus hedonic components of the affective processing of reward.^46^ Consequently, depression in PD may develop due to a pattern of neurodegeneration that disrupts specific striatal subregions disrupting specific frontostriatal circuits which play a crucial role in reward valuation.

The mesolimbic dopamine system has consistently been thought of as a modulator of reward valuation.^24,47^ Several studies have shown a distinction between tonic dopamine signals, which encode reward valuation and effort, and phasic dopamine signals which encode learning.^48–51^ However, the lack of response of reward sensitivity to dopamine in PD depression poses the question: if not dopamine, then what? Neuromodulators including serotonin and noradrenaline have overlapping functions in reward processing and are co-released with dopamine.^52^ Neurodegeneration of the locus coeruleus, the primary site of noradrenaline synthesis in the brain, occur early in the condition and has been associated with apathy and depression in PD.^53,54^ Though noradrenergic function has been predominantly implicated in response vigour and exploratory behaviour, recent studies have shown that pharmacological blockade of noradrenaline leads to a reduction in the use of reward/value information.^55^ Serotenergic dysfunction occurs early in PD^56^ and has been associated with depression^57^, and proposed as a key modulator of reward processing in the brain. In primate studies neuronal recordings from the dorsal raphe nucleus (DRN, one of the main sources of serotonergic neurons) found that firing scaled with the size of prospective reward,^58^ while dietary depletion of serotonin in humans has shown that serotonin selectively modulates reward value during a choice task.^59^ However, our findings do not rule out a role for the dopaminergic system in the aetiology of depression in PD. Dopamine unresponsive reward sensitivity in depressed PD patients could also be explained by dopaminergic medication losing its efficacy due to greater dopaminergic neurodegeneration within key regions involved in reward processing, especially in anhedonic individuals.^16^ We have previously found that motivational symptoms of depression are associated with the degree of dopaminergic degeneration, and that the effects of dopaminergic medications on mood and motivation in PD interact with the degree of striatal dopaminergic neurodegeneration.^27^ In a large longitudinal study of PD patients, monoamine oxidase-B inhibitor treatment improved both depressive and motivation symptoms, but this was attenuated in PD patients with more severe striatal dopaminergic neurodegeneration.^27^ Our results suggest that future studies and novel pharmacological treatment strategies should focus on the interaction between dopamine signaling and other neuromodulators, such as noradrenaline and serotonin, which regulate reward sensitivity.

Reward sensitivity may also be a promising cognitive treatment target for brain stimulation therapies. Recent research has indicated that there are dissociable neural signatures of reward and effort in the brain, with beta oscillations in the basal ganglia tracking subjective effort on a single trial basis and PFC theta oscillations signalling previous trial reward.^60^ The same study goes on to show that deep brain stimulation of the PFC increases reward sensitivity in PD patients,^60^ building on existing evidence that stimulation of this region can selectively modulate willingness to exert effort for reward.^61^ Concurrent DBS targeting of the PFC may be a promising intervention for PD patients who are candidates for DBS therapy for motor symptoms and experience co-morbid treatment resistant depression. However, the recent development of non-invasive neuromodulation using focused ultrasound also enables the potential for targeting deeper brain structure implicated in reward processing and depression such as the ventral striatum in future trials.^62,63^

### Limitations

It remains possible that participants were receiving sub or supratherapeutic dopamine doses that impacted task performance and moderating group differences. However, this is unlikely given the robust group effects, and there were no significant differences in total dopamine dose, disease severity, delay in dopamine, or change in motor symptoms ON and OFF dopamine between groups.

Most participants in both depressed groups were using antidepressants, predominantly selective serotonin reuptake inhibitors (SSRIs) that may have affected task performance. A previous study in healthy participants showed that SSRIs can modulate effort-based decision making reducing effort sensitivity.^64^ However, the same study showed no clear effect on reward sensitivity,^64^ and the majority of trials of SSRIs have shown no significant therapeutic effect on motivational symptoms.^15^ We also found clear differences in in reward sensitivity between depressed and PD depressed groups, suggesting that antidepressant use was not a key moderator of task performance.

Finally, there is the possibility that a confounding factor influenced depressed PD patient performance on the task. As described above we analysed the effect of success during effort exertion to check that probabilistic discounting did not impact offer acceptance, and accounted for this in our modelling, though there were no significant group differences. Reduced concentration or attention, common symptoms in depression, could have impacted task performance. However, we showed specific rather than global changes in decision making, and all groups modulated effort output appropriately.

## Conclusion

This is the first study to investigate the effects of dopamine and depression on effort-based decision-making in PD. We demonstrate that depression in PD is driven by reduced reward sensitivity that is unresponsive to dopamine. This suggests that depression and disruption to effort-based decision-making are not purely related to mesolimbic dopamine function, and other neuromodulatory pathways are likely involved. Our findings indicate that reduced reward sensitivity is a key mechanism and a promising cognitive treatment target for depression in PD that requires non-dopaminergic novel therapies.

## Data availability

Anonymised data are available on request.

## Competing interests

There are no competing interests for authors to disclose.

## Funding

HC is supported by a Wellcome Trust Clinical Training Fellowship, RH is supported by the NIHR UCLH BRC.

## Acknowledgements

We would like to thank all participants who volunteered for this study.

## Supplementary material

**Supplemental figure S1.**
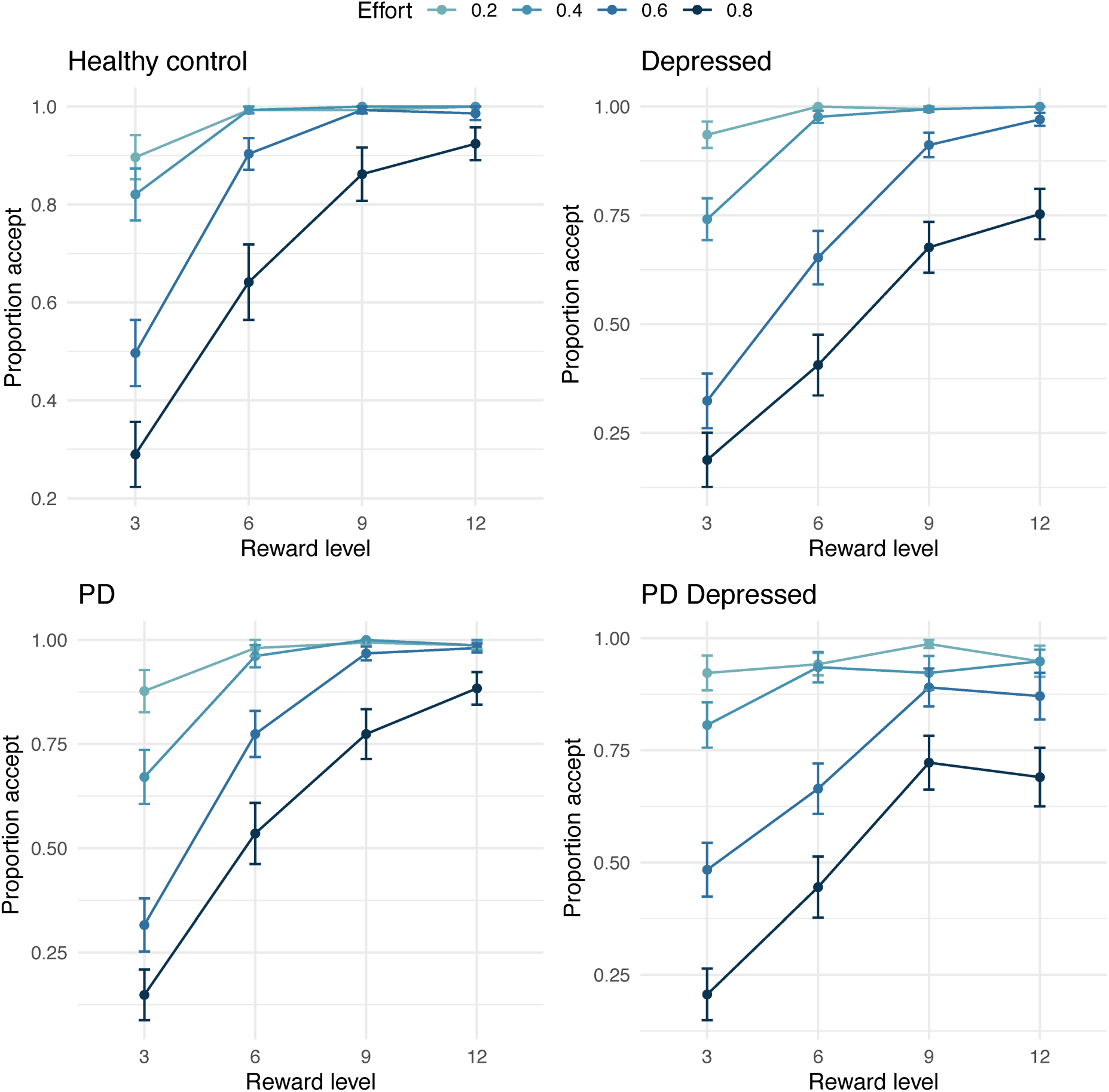
Empirical data showing the change in acceptance rate as reward increases for each effort level by group. (points = means, error bars = standard error)

**Supplemental figure S2.**
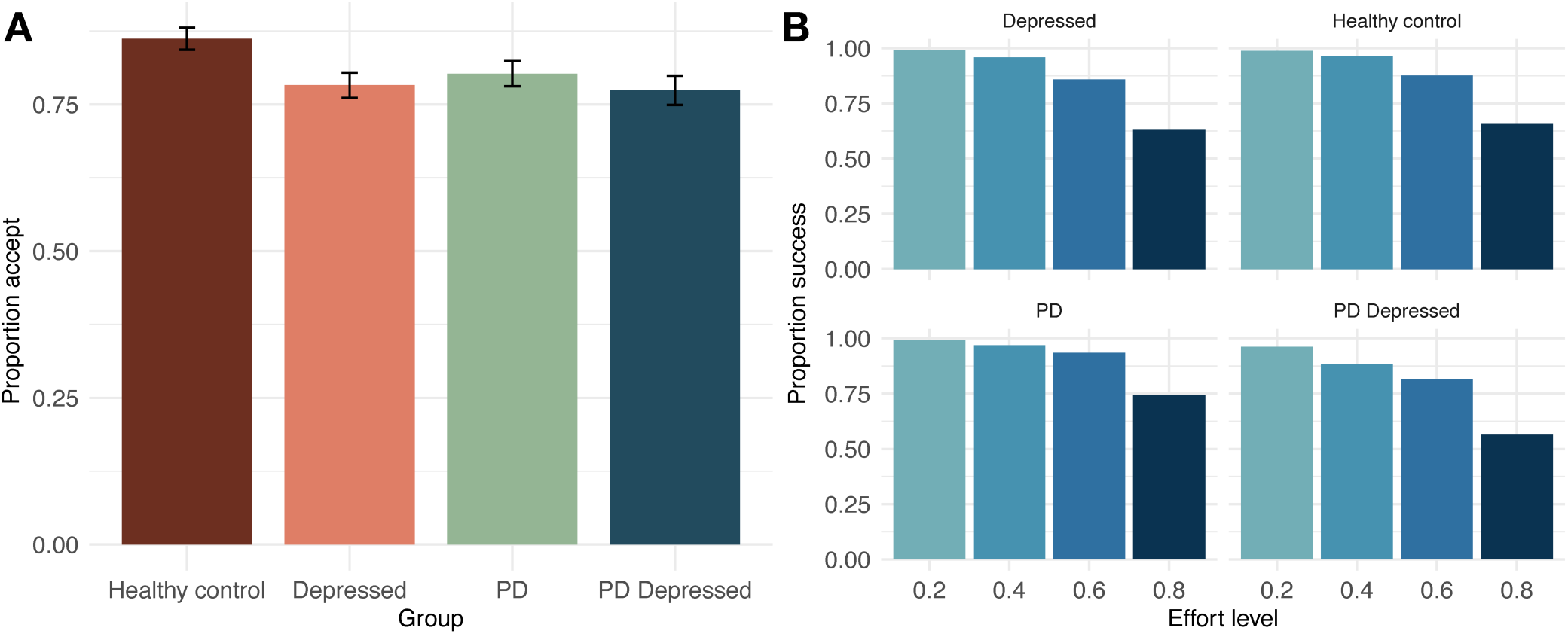
Empirical data showing overall accept rate by group (A) and the change in success rate as effort increases by group. (bars = means, error bars = standard error)

**Supplemental figure S3.**
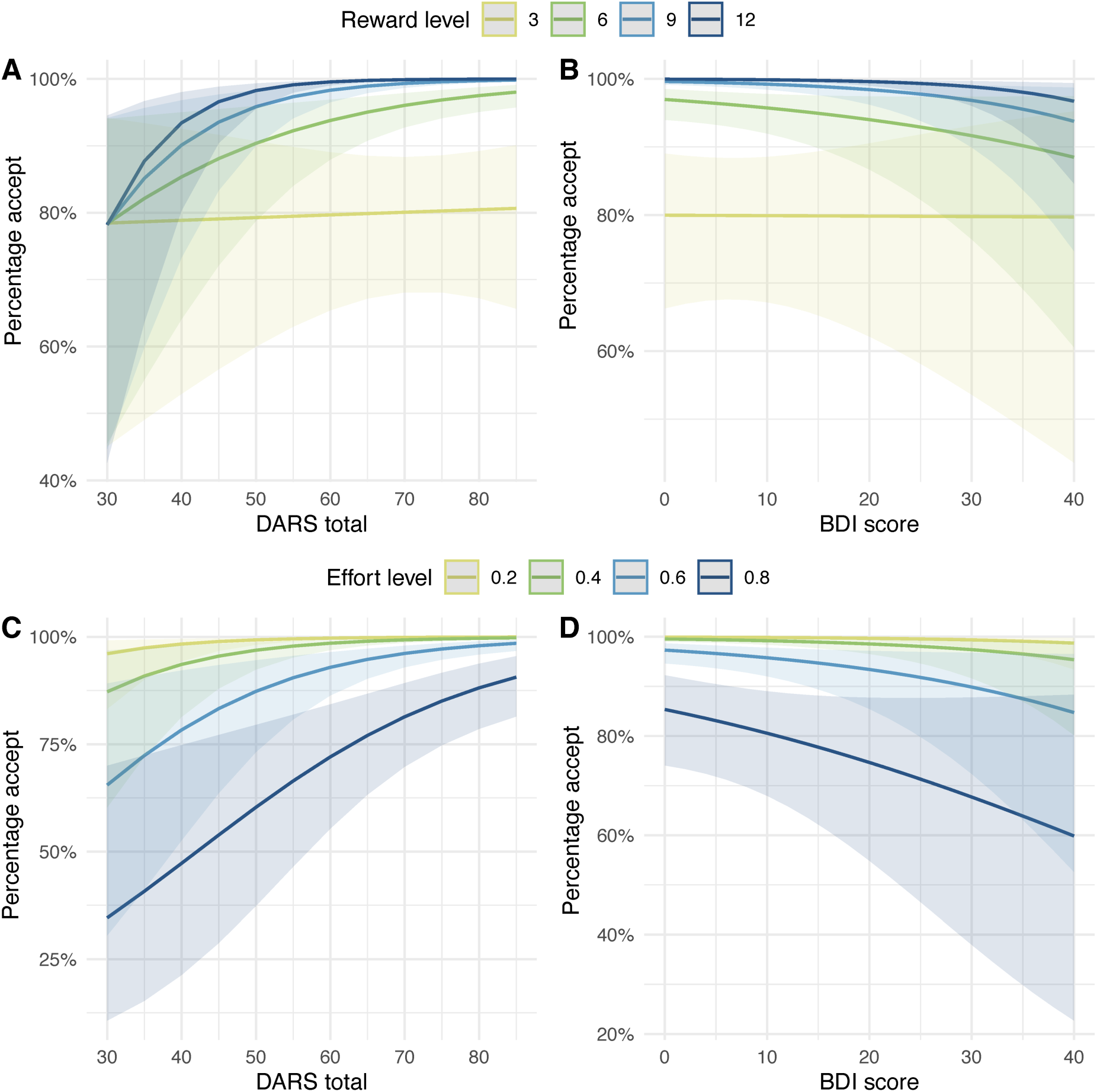
Model predicted plots which shows that participants who were more anhedonic (lower DARS score) or depressed (higher BDI score) were less incentivized by higher rewards to accept an offer (**A** & **B**) and more deterred by higher effort levels. (error bars = 95% confidence interval)

**Supplemental figure S4.**
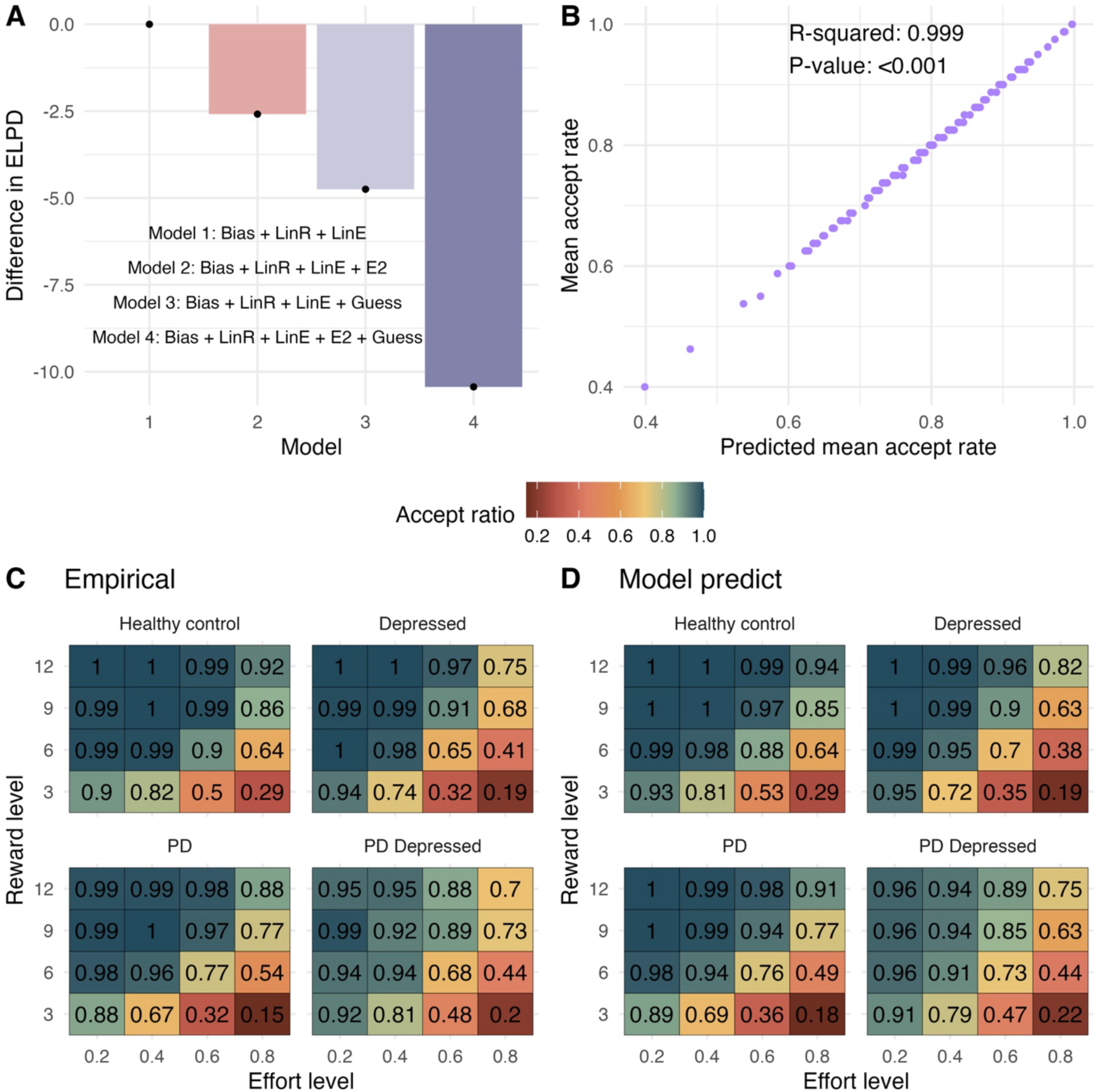
**A.** Model comparison using difference in ELPD (Expected Log Pointwise Predictive Density). ELPD measures the expected log likelihood of new data points under the model, the model with the higher ELPD is more likely to predict new data accurately. Convergence checks were conducted by visualizing trace plots and computing R-hat statistics across MCMC chains. Posterior predictive checks were conducted to ensure model predictions could accurately retrieve behavioural patterns in the original dataset. **B.** Posterior predictive check plot of empirical mean individual accept rate and predicted individual mean accept rate showing almost perfect recovery of individual differences in acceptance rates. **C** & **D**. Change in each group acceptance as both reward and effort increases seen in empirical (**C**) and model predicted data (**D**), demonstrating qualitatively similar results.

**Supplemental figure S5.**
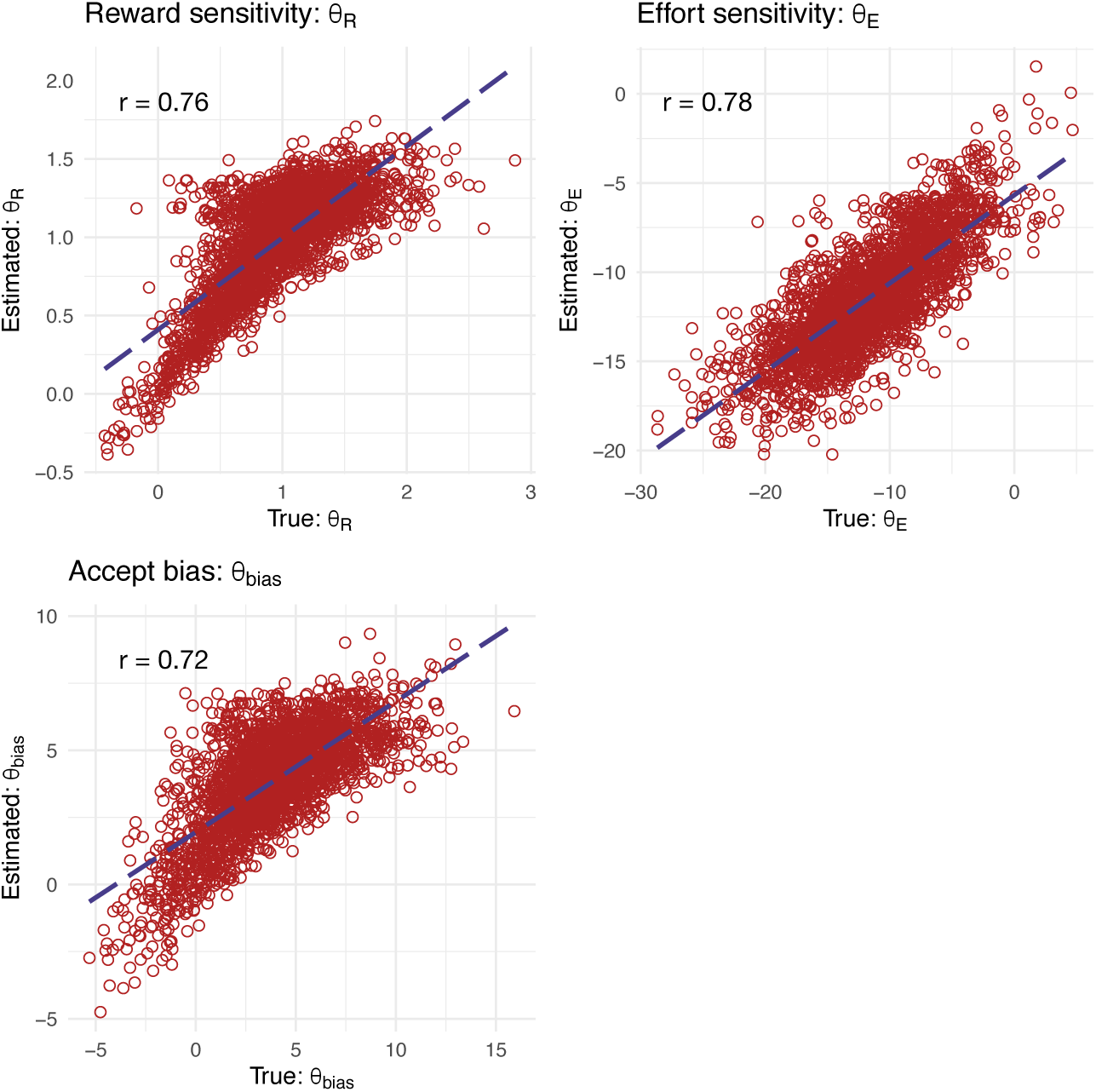
Parameter recovery for the winning model demonstrated excellent recovery with Pearson’s r between data generating and recovered parameters ≈ 0.72 – 0.78.

**Supplemental figure S6.**
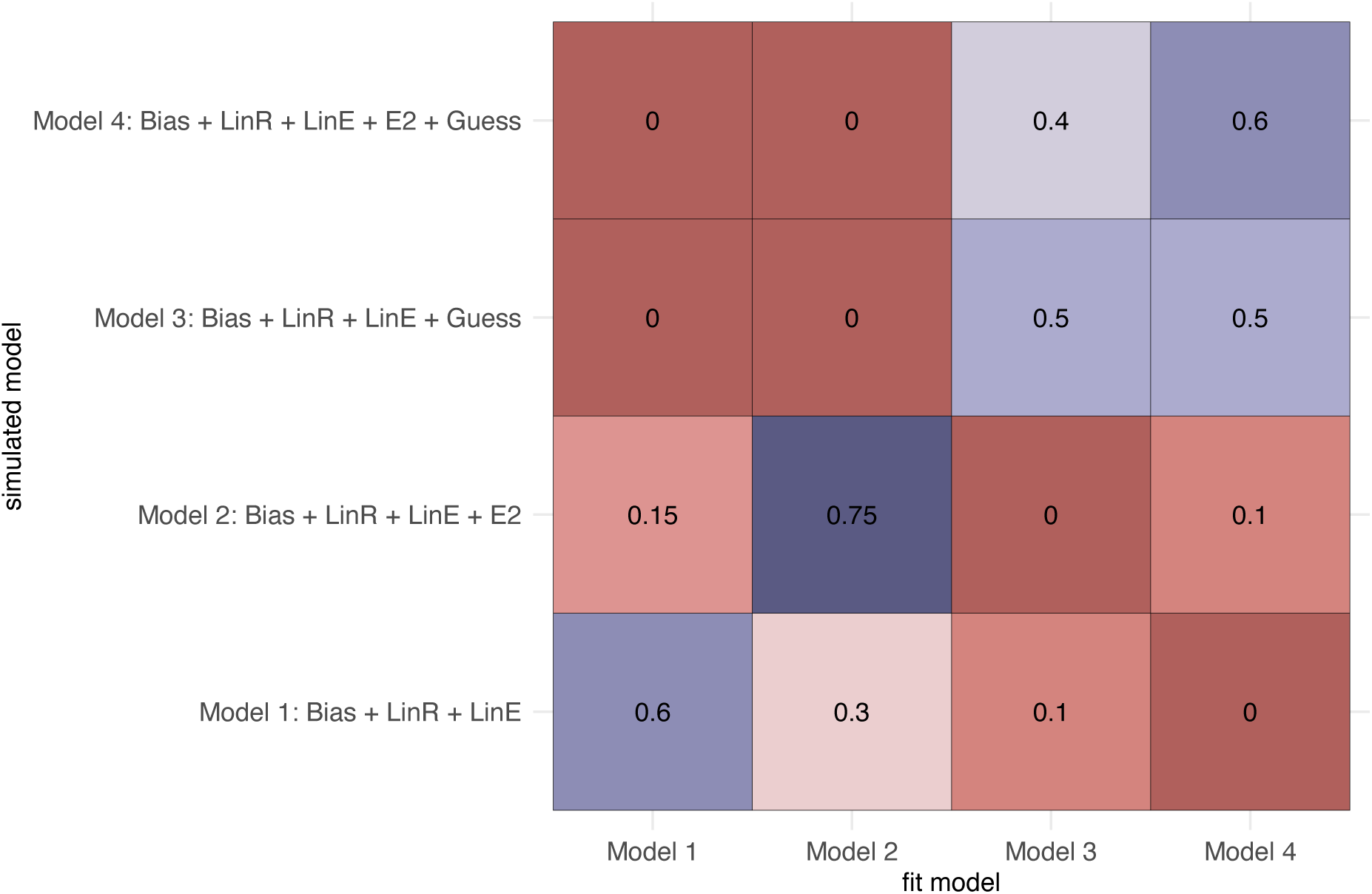
Confusion matrix showing the effect of prior parameter distributions on model recovery. Numbers denote the probability that data generated with model X (simulated model) are best fit by model Y (fit model)

**Supplemental Figure 7.**
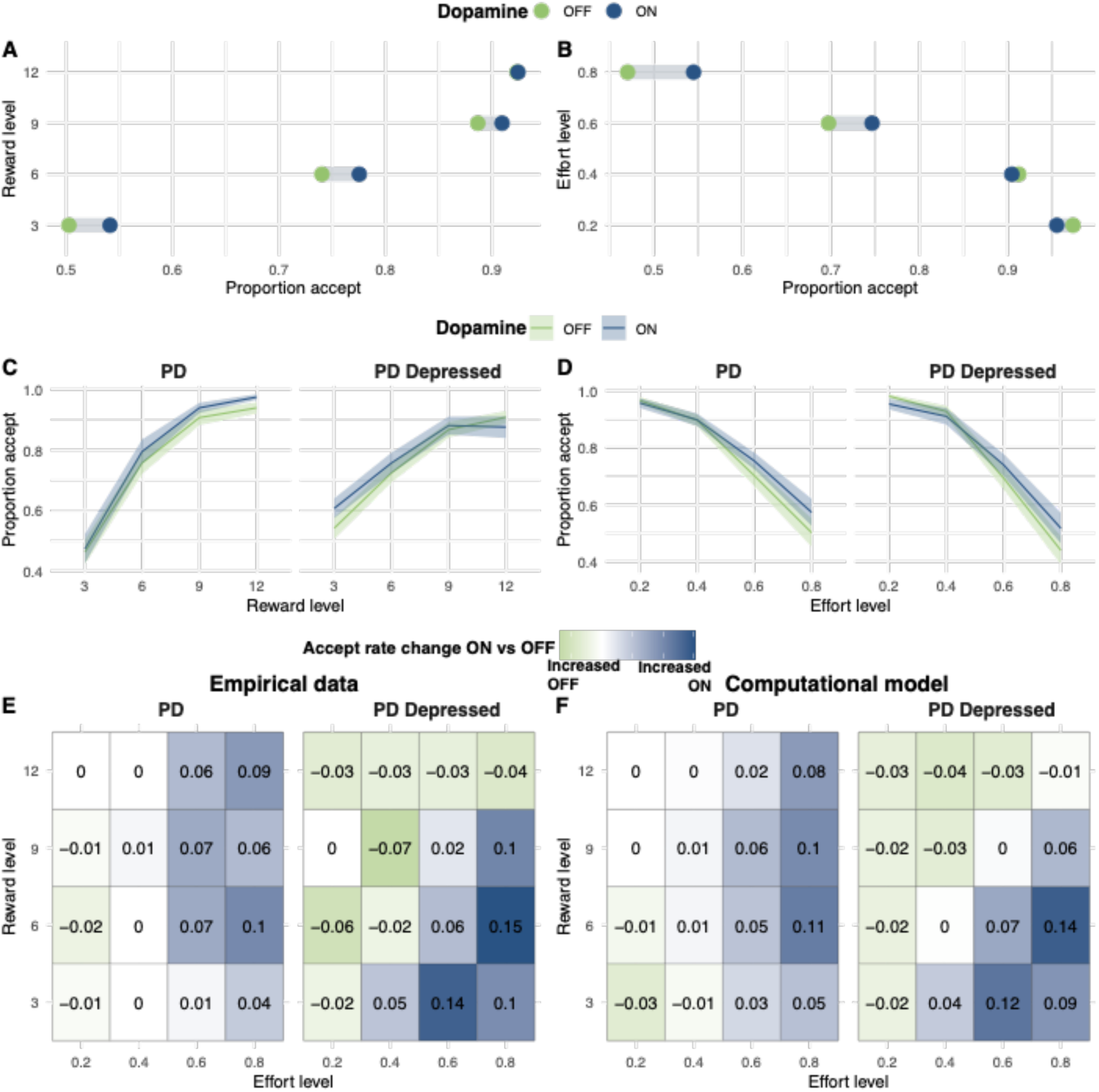
Empirical data showing mean change in acceptance rate ON and OFF dopamine medication as reward (**A**) and effort (**B**) increase, across both PD groups. Acceptance rate for PD and PD depressed groups ON and OFF dopamine medication as reward (**C**) and (**D**) effort increase. Change in acceptance rate with dopamine medication (ON minus OFF) for PD and PD depressed groups as reward and effort increase in (**E**) empirical data and (**F**) simulated data from the Bayesian computational model.

**Supplemental Table 1.**
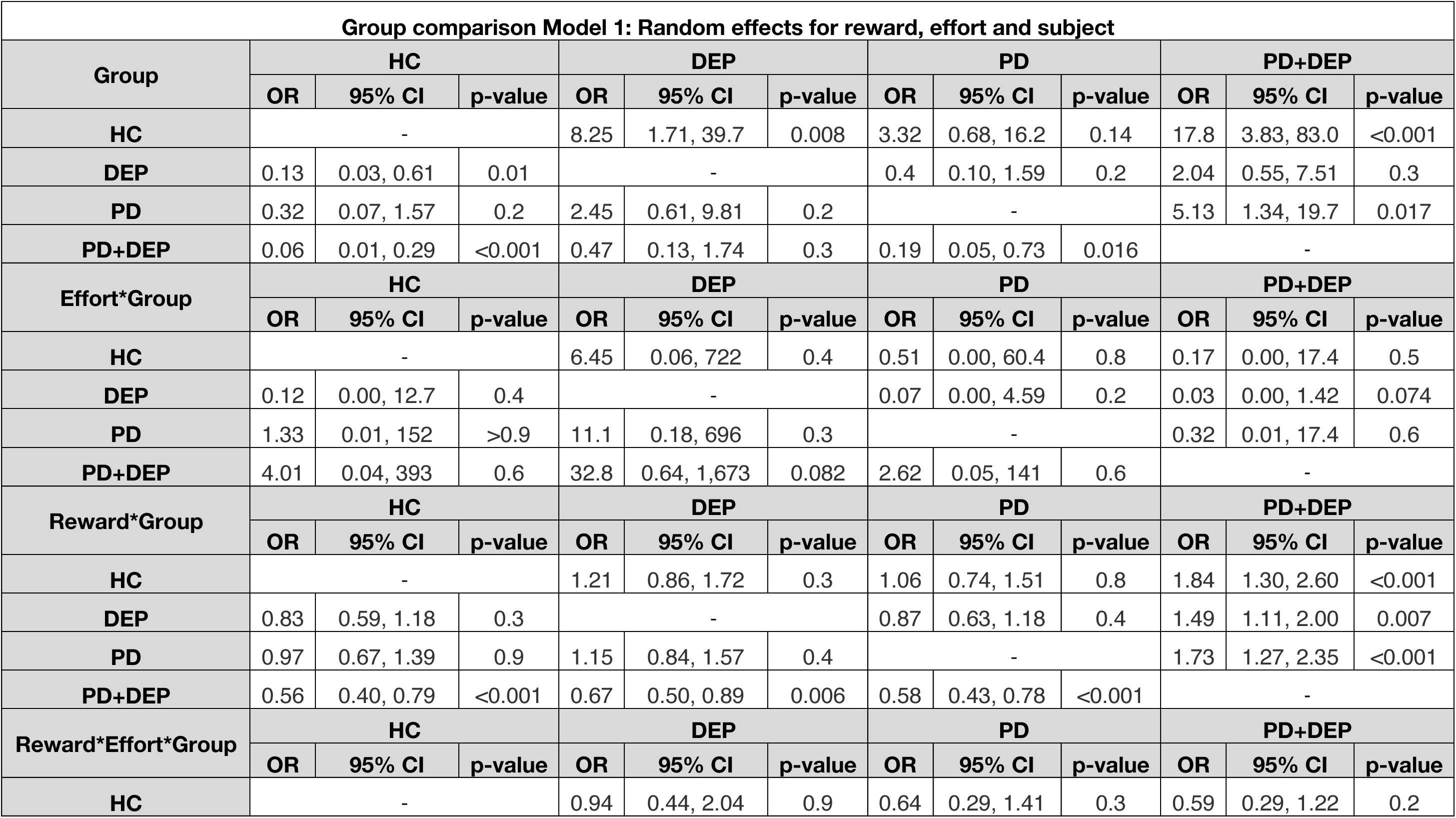
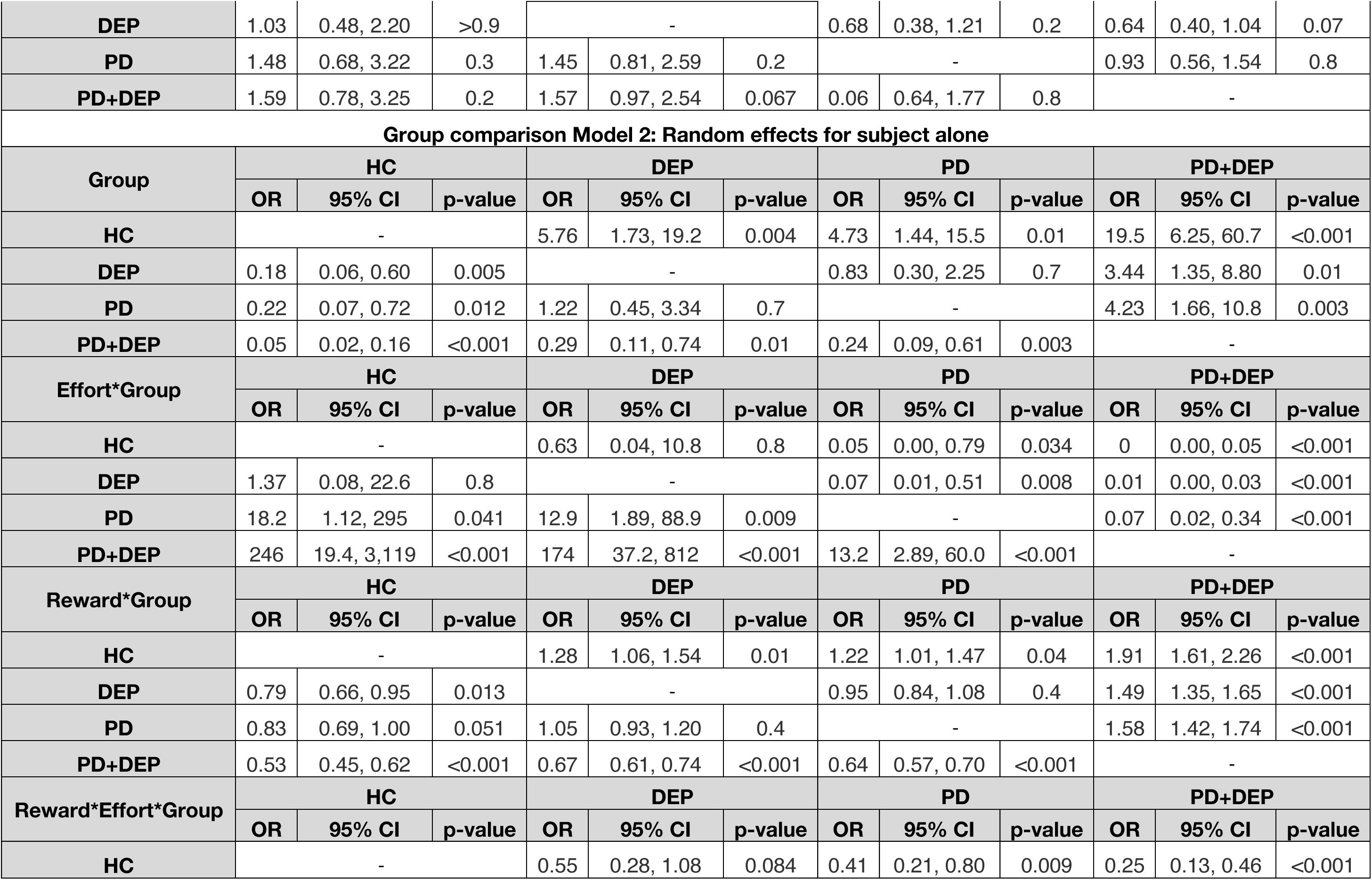
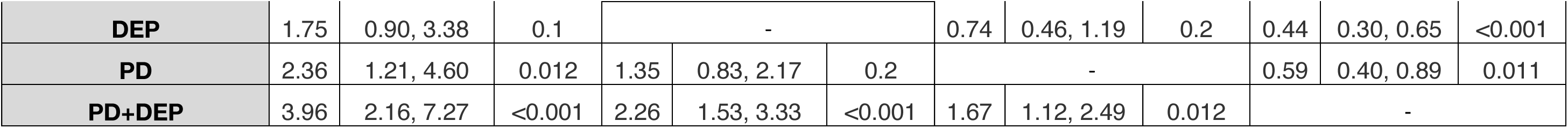
Mixed effects models of group comparison with and without the addition of random effects for reward and effort to subject alone.

| Group comparison Model 1: Random effects for reward, effort and subject |  |  |  |  |  |  |  |  |  |  |  |  |
| --- | --- | --- | --- | --- | --- | --- | --- | --- | --- | --- | --- | --- |
| Group | HC |  |  | DEP |  |  | PD |  |  | PD+DEP |  |  |
|  | OR | 95% CI | p-value | OR | 95% CI | p-value | OR | 95% CI | p-value | OR | 95% CI | p-value |
| HC | - |  |  | 8.25 | 1.71, 39.7 | 0.008 | 3.32 | 0.68, 16.2 | 0.14 | 17.8 | 3.83, 83.0 | <0.001 |
| DEP | 0.13 | 0.03, 0.61 | 0.01 | - |  |  | 0.4 | 0.10, 1.59 | 0.2 | 2.04 | 0.55, 7.51 | 0.3 |
| PD | 0.32 | 0.07, 1.57 | 0.2 | 2.45 | 0.61, 9.81 | 0.2 | - |  |  | 5.13 | 1.34, 19.7 | 0.017 |
| PD+DEP | 0.06 | 0.01, 0.29 | <0.001 | 0.47 | 0.13, 1.74 | 0.3 | 0.19 | 0.05, 0.73 | 0.016 | - |  |  |
| Effort*Group | HC |  |  | DEP |  |  | PD |  |  | PD+DEP |  |  |
|  | OR | 95% CI | p-value | OR | 95% CI | p-value | OR | 95% CI | p-value | OR | 95% CI | p-value |
| HC | - |  |  | 6.45 | 0.06, 722 | 0.4 | 0.51 | 0.00, 60.4 | 0.8 | 0.17 | 0.00, 17.4 | 0.5 |
| DEP | 0.12 | 0.00, 12.7 | 0.4 | - |  |  | 0.07 | 0.00, 4.59 | 0.2 | 0.03 | 0.00, 1.42 | 0.074 |
| PD | 1.33 | 0.01, 152 | >0.9 | 11.1 | 0.18, 696 | 0.3 | - |  |  | 0.32 | 0.01, 17.4 | 0.6 |
| PD+DEP | 4.01 | 0.04, 393 | 0.6 | 32.8 | 0.64, 1,673 | 0.082 | 2.62 | 0.05, 141 | 0.6 | - |  |  |
| Reward*Group | HC |  |  | DEP |  |  | PD |  |  | PD+DEP |  |  |
|  | OR | 95% CI | p-value | OR | 95% CI | p-value | OR | 95% CI | p-value | OR | 95% CI | p-value |
| HC | - |  |  | 1.21 | 0.86, 1.72 | 0.3 | 1.06 | 0.74, 1.51 | 0.8 | 1.84 | 1.30, 2.60 | <0.001 |
| DEP | 0.83 | 0.59, 1.18 | 0.3 | - |  |  | 0.87 | 0.63, 1.18 | 0.4 | 1.49 | 1.11, 2.00 | 0.007 |
| PD | 0.97 | 0.67, 1.39 | 0.9 | 1.15 | 0.84, 1.57 | 0.4 | - |  |  | 1.73 | 1.27, 2.35 | <0.001 |
| PD+DEP | 0.56 | 0.40, 0.79 | <0.001 | 0.67 | 0.50, 0.89 | 0.006 | 0.58 | 0.43, 0.78 | <0.001 | - |  |  |
| Reward*Effort*Group | HC |  |  | DEP |  |  | PD |  |  | PD+DEP |  |  |
|  | OR | 95% CI | p-value | OR | 95% CI | p-value | OR | 95% CI | p-value | OR | 95% CI | p-value |
| HC | - |  |  | 0.94 | 0.44, 2.04 | 0.9 | 0.64 | 0.29, 1.41 | 0.3 | 0.59 | 0.29, 1.22 | 0.2 |
| DEP | 1.03 | 0.48, 2.20 | >0.9 | - |  |  | 0.68 | 0.38, 1.21 | 0.2 | 0.64 | 0.40, 1.04 | 0.07 |
| PD | 1.48 | 0.68, 3.22 | 0.3 | 1.45 | 0.81, 2.59 | 0.2 | - |  |  | 0.93 | 0.56, 1.54 | 0.8 |
| PD+DEP | 1.59 | 0.78, 3.25 | 0.2 | 1.57 | 0.97, 2.54 | 0.067 | 0.06 | 0.64, 1.77 | 0.8 | - |  |  |
| Group comparison Model 2: Random effects for subject alone |  |  |  |  |  |  |  |  |  |  |  |  |
| Group | HC |  |  | DEP |  |  | PD |  |  | PD+DEP |  |  |
|  | OR | 95% CI | p-value | OR | 95% CI | p-value | OR | 95% CI | p-value | OR | 95% CI | p-value |
| HC | - |  |  | 5.76 | 1.73, 19.2 | 0.004 | 4.73 | 1.44, 15.5 | 0.01 | 19.5 | 6.25, 60.7 | <0.001 |
| DEP | 0.18 | 0.06, 0.60 | 0.005 | - |  |  | 0.83 | 0.30, 2.25 | 0.7 | 3.44 | 1.35, 8.80 | 0.01 |
| PD | 0.22 | 0.07, 0.72 | 0.012 | 1.22 | 0.45, 3.34 | 0.7 | - |  |  | 4.23 | 1.66, 10.8 | 0.003 |
| PD+DEP | 0.05 | 0.02, 0.16 | <0.001 | 0.29 | 0.11, 0.74 | 0.01 | 0.24 | 0.09, 0.61 | 0.003 | - |  |  |
| Effort*Group | HC |  |  | DEP |  |  | PD |  |  | PD+DEP |  |  |
|  | OR | 95% CI | p-value | OR | 95% CI | p-value | OR | 95% CI | p-value | OR | 95% CI | p-value |
| HC | - |  |  | 0.63 | 0.04, 10.8 | 0.8 | 0.05 | 0.00, 0.79 | 0.034 | 0 | 0.00, 0.05 | <0.001 |
| DEP | 1.37 | 0.08, 22.6 | 0.8 | - |  |  | 0.07 | 0.01, 0.51 | 0.008 | 0.01 | 0.00, 0.03 | <0.001 |
| PD | 18.2 | 1.12, 295 | 0.041 | 12.9 | 1.89, 88.9 | 0.009 | - |  |  | 0.07 | 0.02, 0.34 | <0.001 |
| PD+DEP | 246 | 19.4, 3,119 | <0.001 | 174 | 37.2, 812 | <0.001 | 13.2 | 2.89, 60.0 | <0.001 | - |  |  |
| Reward*Group | HC |  |  | DEP |  |  | PD |  |  | PD+DEP |  |  |
|  | OR | 95% CI | p-value | OR | 95% CI | p-value | OR | 95% CI | p-value | OR | 95% CI | p-value |
| HC | - |  |  | 1.28 | 1.06, 1.54 | 0.01 | 1.22 | 1.01, 1.47 | 0.04 | 1.91 | 1.61, 2.26 | <0.001 |
| DEP | 0.79 | 0.66, 0.95 | 0.013 | - |  |  | 0.95 | 0.84, 1.08 | 0.4 | 1.49 | 1.35, 1.65 | <0.001 |
| PD | 0.83 | 0.69, 1.00 | 0.051 | 1.05 | 0.93, 1.20 | 0.4 | - |  |  | 1.58 | 1.42, 1.74 | <0.001 |
| PD+DEP | 0.53 | 0.45, 0.62 | <0.001 | 0.67 | 0.61, 0.74 | <0.001 | 0.64 | 0.57, 0.70 | <0.001 | - |  |  |
| Reward*Effort*Group | HC |  |  | DEP |  |  | PD |  |  | PD+DEP |  |  |
|  | OR | 95% CI | p-value | OR | 95% CI | p-value | OR | 95% CI | p-value | OR | 95% CI | p-value |
| HC | - |  |  | 0.55 | 0.28, 1.08 | 0.084 | 0.41 | 0.21, 0.80 | 0.009 | 0.25 | 0.13, 0.46 | <0.001 |
| **DEP** | 1.75 | 0.90, 3.38 | 0.1 | - |  |  | 0.74 | 0.46, 1.19 | 0.2 | 0.44 | 0.30, 0.65 | <0.001 |
| **PD** | 2.36 | 1.21, 4.60 | 0.012 | 1.35 | 0.83, 2.17 | 0.2 | - |  |  | 0.59 | 0.40, 0.89 | 0.011 |
| **PD+DEP** | 3.96 | 2.16, 7.27 | <0.001 | 2.26 | 1.53, 3.33 | <0.001 | 1.67 | 1.12, 2.49 | 0.012 | - |  |  |

## Notes

### Competing Interest Statement

The authors have declared no competing interest.

### Summary of Updates

Additional analysis, minor corrections and repeat modelling.

